# Effects of a 50-Hz electric field on sleep quality and life span mediated by ultraviolet (UV)-A/blue light photoreceptor CRYPTOCHROME in *Drosophila melanogaster*

**DOI:** 10.1101/2020.09.23.310862

**Authors:** Haruhisa Kawasaki, Hideyuki Okano, Takaki Nedachi, Yuzo Nakagawa-Yagi, Akikuni Hara, Norio Ishida

## Abstract

Although electric fields (EF) exert beneficial effects on animal wound healing and differentiation, the molecular mechanisms of these effects have remained unclear for years. Therefore, we aimed to elucidate the molecular mechanisms underlying these effects in *Drosophila melanogaster* as a genetic animal model. The sleep quality of wild-type (WT) flies was improved by exposure to a 50-Hz (35 kv/m) constant electric field during the daytime, but not during the night. This effect was undetectable in *Cryptochrome* mutant (*Cry^b^*) flies. Exposure to a 50-Hz electric field under low nutrient conditions elongated the lifespan of male and female WT flies by ~18%, but not of three diferrent *Cry* mutants and *Cry* RNAi strains. Metabolome analysis indicated that the adenosine triphosphate (ATP) content was 5-fold higher in intact WT than *Cry* gene mutant strains exposed to an electric field.

A putative magnetoreceptor protein and UV-A/blue light photoreceptor, CRYPTOCHROME (CRY) is involved in electric field receptors in animals. The present findings constitute hitherto unknown genetic evidence of a CRY-based system that is electric-field sensitive in animals.

## Introduction

Electric fields (EF) influence various behaviors, including embryogenesis, wound healing, polarity, differentiation, and motility in plants and animals (1–3). However, how electrical cues are received and translated into a cellular response in most life systems is poorly understood.

High voltage electric fields (EF) are effective for treating stiff shoulders, headaches, insomnia and chronic constipation (4). The Japanese Ministry of Health, Labor and Welfare has approved a device that that delivers high-voltage electrical potentials (HELP) to the body. Although EF therapy was discovered ~60 years ago, the molecular mechanisms of its benefits have remained unknown (5), despite investigations involving humans and mice (6). Therefore *Drosophila* melanogaster was the experimental model in the present study.

Several studies have found that animals and plants detect magnetic fields through CRY proteins. Magnetic fields affect the behavior of cockroaches (7), flies (8–10), butterflies (11), birds (12, 13) and humans (14). The *Cryptochrome (Cry*) gene plays a critical role in magnetic detection among diverse species. CRY protein forms complexes with MagR protein in the retina of immigrant birds indicating that they can recognize geomagnetics (8).

The present findings showed that an EF exposure of 50 Hz can improve sleep quality and extend the lifetime in WT flies and that *Cry* is essential for the electric field receptor system in *Drosophila*.

## Materials and methods

### Flies

The *Drosophila* strains Oregon R, *Cry^b^, Cry^01^, Cry^03^* (a gift from Dr.Paul Hardin, Texas A&M), *w^1118^; P{GD738}v7239* from the Vienna *Drosophila* Resource Center (VDRC, www.vdrc.at) and *Gal4-elav* were maintained as described (15, 16). The transgenic strain *w^1118^; P{GD738}v7239* contains upstream regulatory sequences (UAS) to drive the RNAi for the *Cry* gene.

### Electric field exposure

We created a uniform EF by transforming a 50-Hz alternating current at 35 kV/m using a Healthtron HEF-P3500 (Hakuju Institute for Health Science Co., Ltd., Tokyo, Japan) to deliver EF. Flies and medium were placed in vials or tubes and placed 3-5 cm apart between the electrodes of the Healthtron for EF exposure.

### Assays of sleep behavior

Male flies were individually placed in glass tubes (inner diameter, 3 mm) containing 5% sucrose and 2% agarose and exposed to EF. The tubes were then transferred to a Drosophila Activity Monitoring (DAM) system (TriKinetics Inc., Waltham, MA, USA) and locomotor activity was measured in 1-min bins. Sleep was defined as > 5 min of consolidated inactivity (17, 18)and > 60 min for more detailed analysis.

### Metabolome analysis

Two days after eclosion, male Oregon R or *Cry^b^* flies were exposed to EF for 48 h, then stored (30 - 40 mg batches) frozen in liquid nitrogen. Metabolites were extracted and metabolomes were measured at Human Metabolome Technologies Inc. (HMT, Tsuruoka, Japan. For CE-TOFMS analysis, 1,500 μL of 50% acetonitrile (v/v) was added to *Drosophila* samples and blended under cooling using a BMS-M10N21 homogenizer (Bio Medical Science Inc., Tokyo, Japan) at 1,500 rpm, 120 sec × 5). Homogenates were separated by centrifugation at 2,300 × g for 5 min at 4°C and the upper aqueous layer was centrifugally passed through Millipore 5 kDa cut-of filter (Ultrafree MC-PLHCC, HMT; Millipore Sigma Co., Ltd., Burlington, MA, USA) at 9,100 × g for 120 min at 4°C to remove macromolecules. The filtrate was then concentrated by centrifugation and reconstituted in 100 μL of Milli-Q water before CE-TOFMS analysis.

Cationic metabolites were analyzed using a fused silica capillary (50 μm i.d. × 80 cm), with the cation electrophoresis buffer H3301-1001 (Human Metabolome Technologies) as the electrolyte. Samples were injected at a pressure of 50 mbar for 10 s, and the applied voltage was 27 kV. Samples were assessed by electrospray ionization-mass spectrometry (ESI-MS) in the positive ion mode, at a capillary voltage of 4000 V. Spectra were obtained between m/z 50 and 1,000.

Anionic metabolites were analyzed using a fused silica capillary (50 μm i.d. × 80 cm), with the anionic electrophoresis buffer H3302-1021 (Human Metabolome Technologies) as the electrolyte. Samples were injected at a pressure of 50 mbar for 25 s, and the applied voltage was 30 kV. Samples were assessed by ESI-MS in the negative ion mode, and the capillary voltage was 3,500 V. Spectra were obtained between m/z 50 and 1,000.

### Lifespan measurements

Lifespan under low nutrient conditions (low nutrient resistance) was assessed by placing male and female flies (n = 30 each) in vials containing 5 mL of 1% agarose and 5% glucose at 25°C under LD 12:12 cycles. Dead flies were counted every 2–3 days. Lifespan under starvation in 1% agarose was similarly assayed.

## Results

### Daytime exposure to an electric field improves sleep quality

We assayed sleep in *Drosophila* to determine the effects of a 50-Hz EF on health. The sleep quality of WT flies exposed to a 12-h EF during the day (ZT 0-12) or night (ZT 13-24) differed (Fig. 1a). Exposure to EF during the day decreased sleep bout number during the night, but increased the sleep bout length and total sleep in WT flies (Fig. 1b). These results suggest that sleep fragmentation was avoided after daytime EF exposure.

**Fig. 1.**
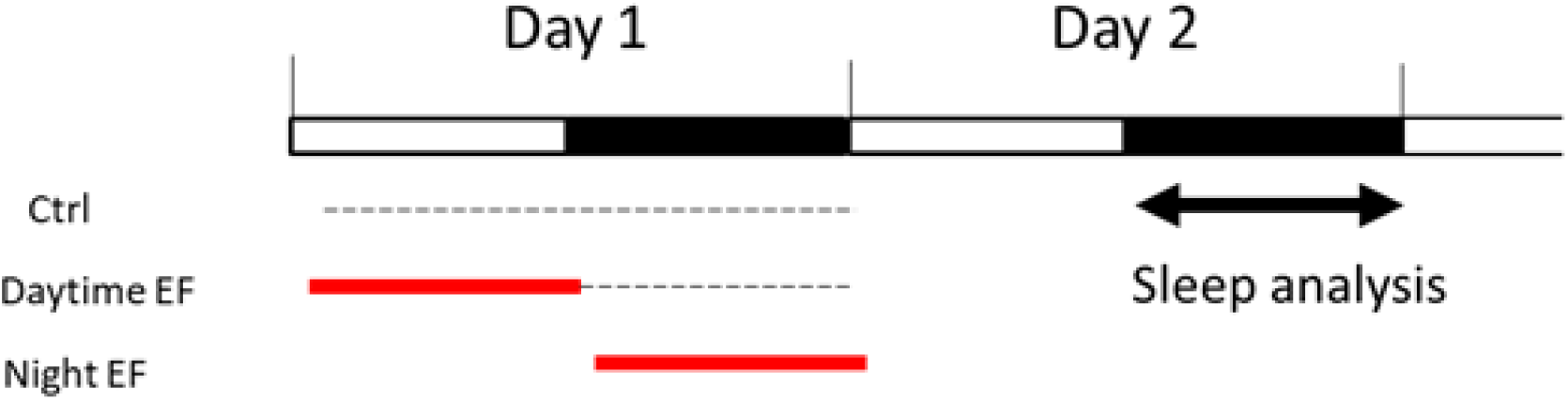
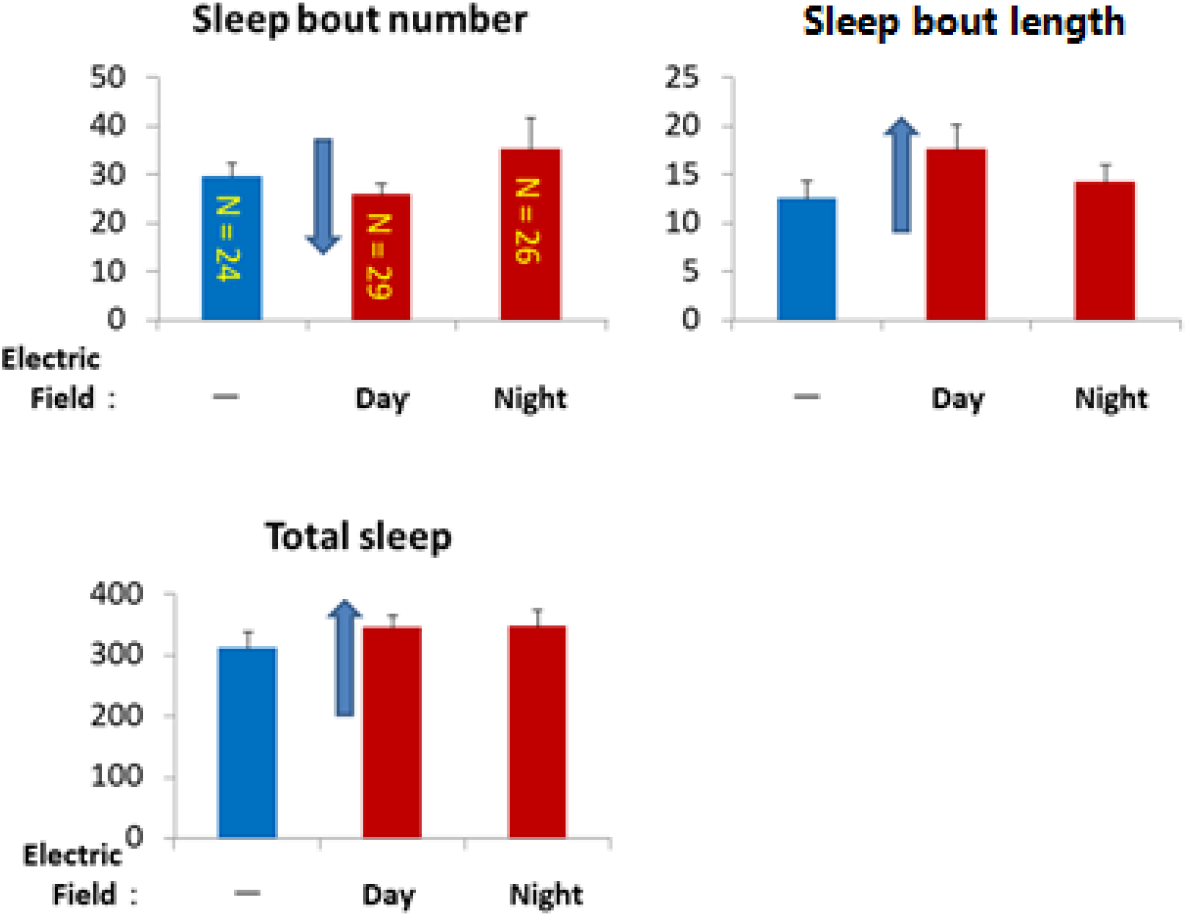
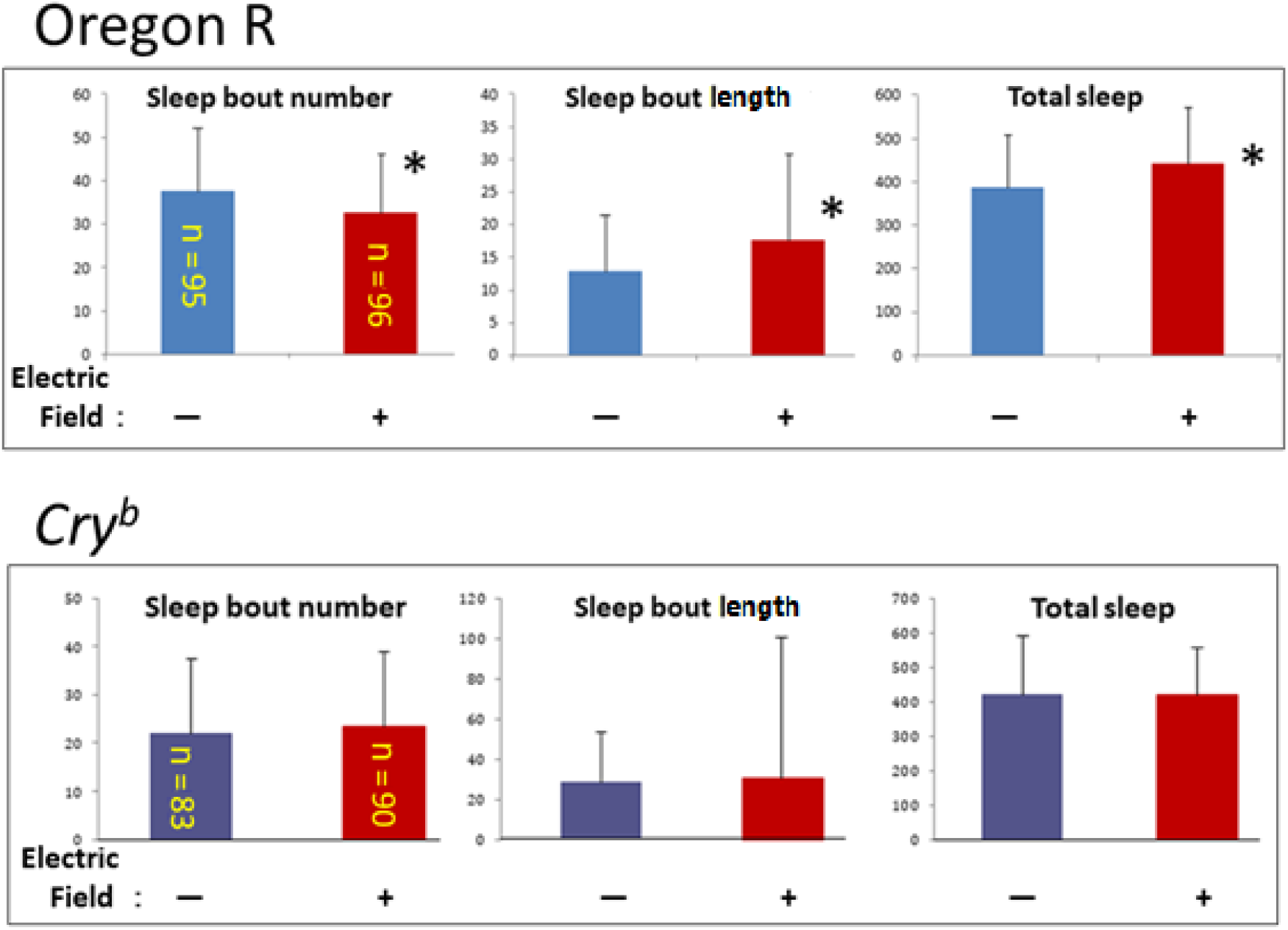
Sleep quality was improved by daytime EF exposure. a. Assessment of EF effects on sleep in WT *Drosophila* (Oregon R). Nighttime sleep was analyzed after daytime (DEF) or nighttime (NEF) exposure to EF for 12 h. b. Comparison of DEF and NEF exposure (shown above) on nighttime sleep between WT *Drosophila* on day 2. Sleep bouts, duration and total sleep were determined in control flies (-). Daytime EF exposure decreased sleep bout number, and increased sleep bout length and total sleep time in Oregon-R. c. Effects of daytime EF exposure on nighttime sleep in clock mutant *Cry^b^* and WT (Oregon R) flies. Two days after EF exposure, we determined sleep analysis as sum of four independent experiments. Daytime EF decreased sleep bout number, and increased bout length and total sleep in WT, but did not significantly affect these in *Cry* mutant flies. EF, electric field, WT, wild type.

We explored this possibility by evaluating sleep in a large number of WT and *Cryptochrome* mutant flies. The results were the same (avoidance of sleep fragmentation) in the WT but not in a *Cryptochrome* mutant, *Cry^b^* (Fig. 1c). We concluded that daytime exposure to a 50-Hz EF improved sleep quality via regulating the *Cryptochrome* gene.

### Exposure to EF prolongs life span under low nutrients

We maintained flies under low nutritional status then exposed them to EF to determine its effects on lifespan. The lifespan of WT flies exposed to EF was significantly increased ~18% compared with that of sham-treated control flies (Fig. 2a). These results showed that exposure to an EF increases the lifespan *of* Drosophila under low nutritional conditions.

**Fig. 2.**
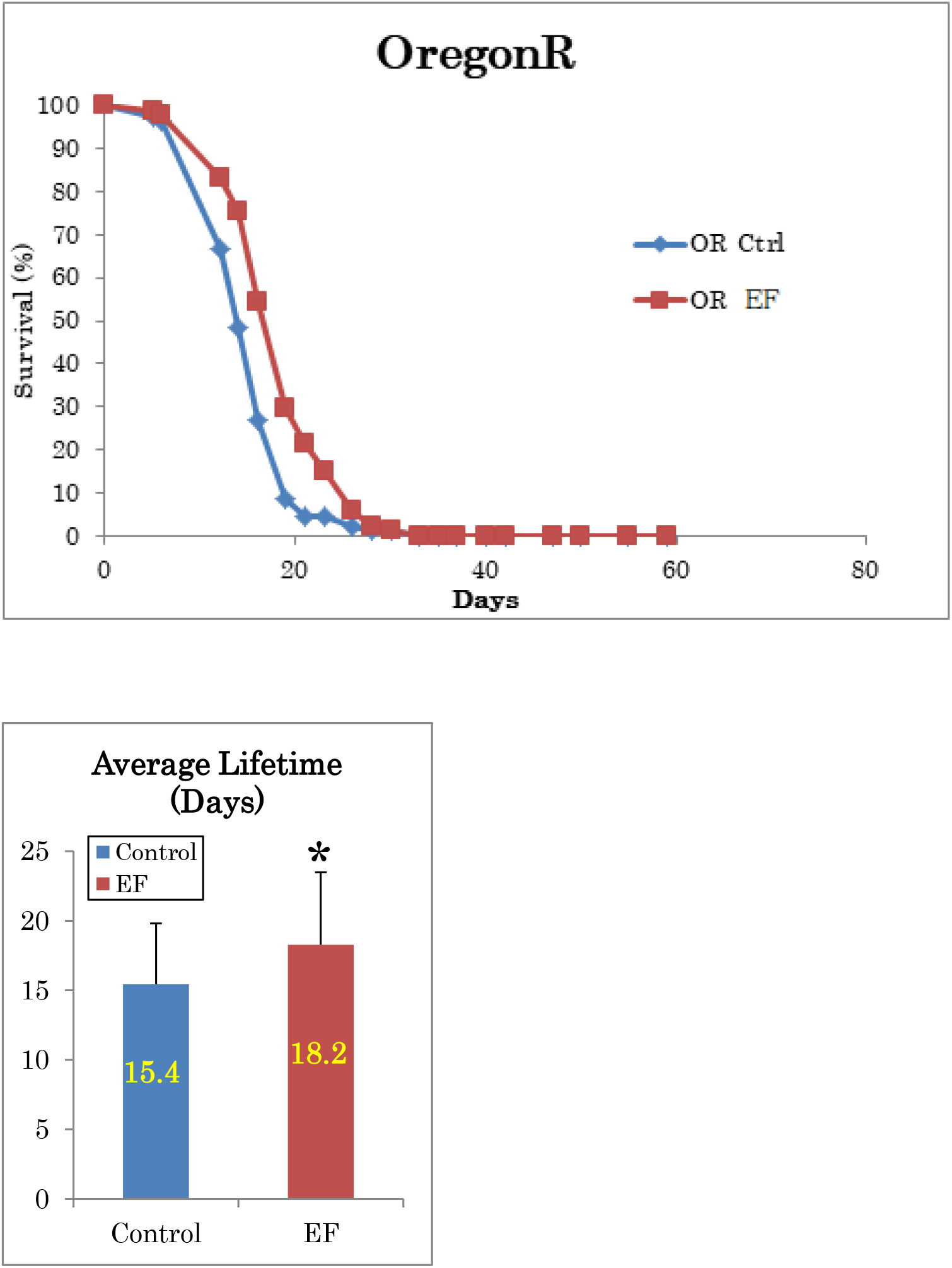
Exposure to EF prolonged lifespan of WT flies maintained under low nutrient food conditions. a. Flies under low nutritional status were exposed to 50-Hz EF. b. Graph shows average lifespans. EF, electric field, WT, wild type. b, The lifespans of EF exposed WT was increased ~18% than shame control.

### Exposure EF increases lifespan of *Drosophila* under starvation

We compared the lifespans of starved male and female WT flies exposed to EF to determine whether the longevity effect would persist. The lifespan of both sexes were significantly increased by ~20% in these flies compared with sham-treated flies (Fig. 3a). These results showed that EF increased the lifespan of WT flies under starvation and in low nutritional conditions.

**Fig. 3.**
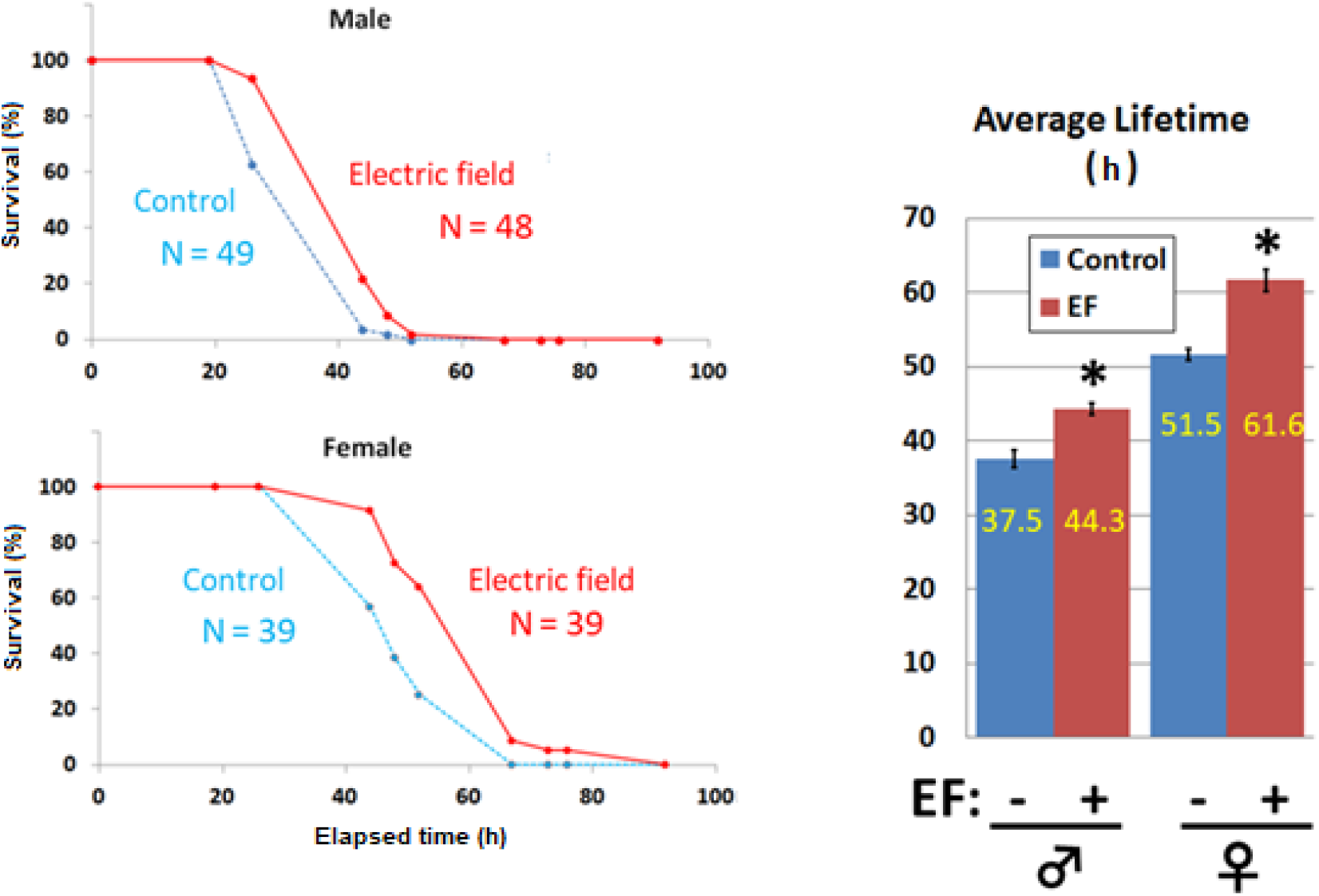
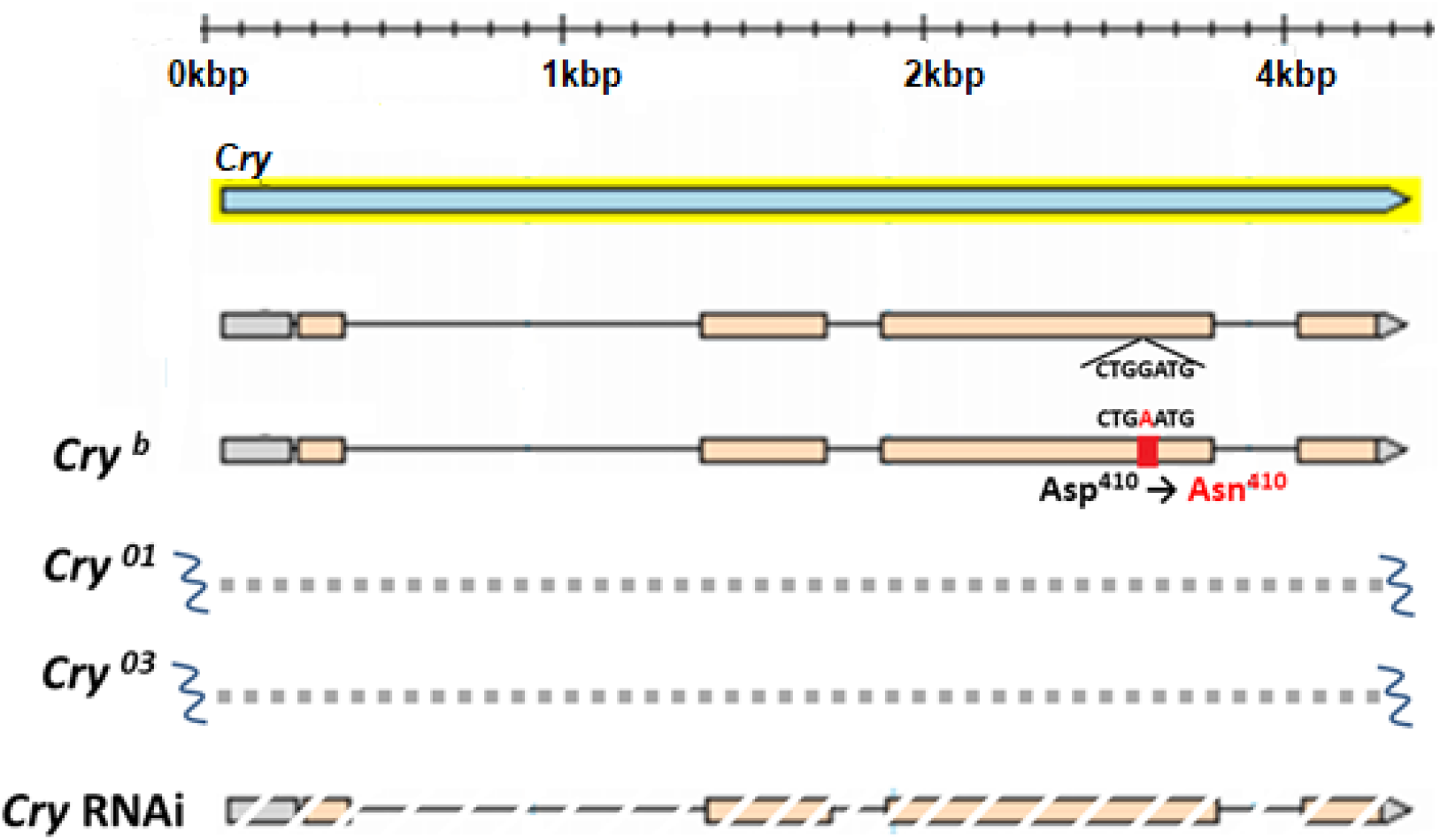
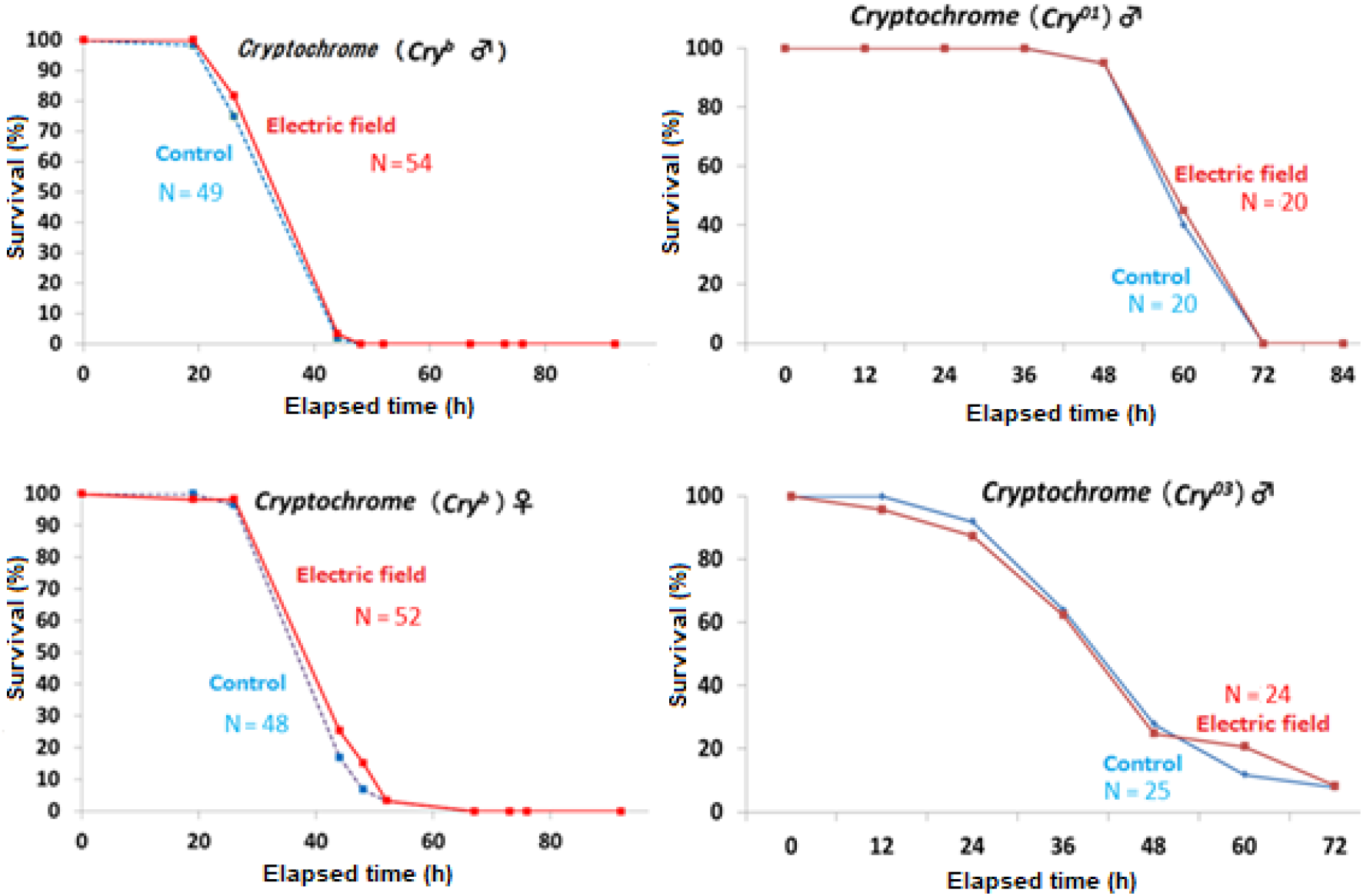
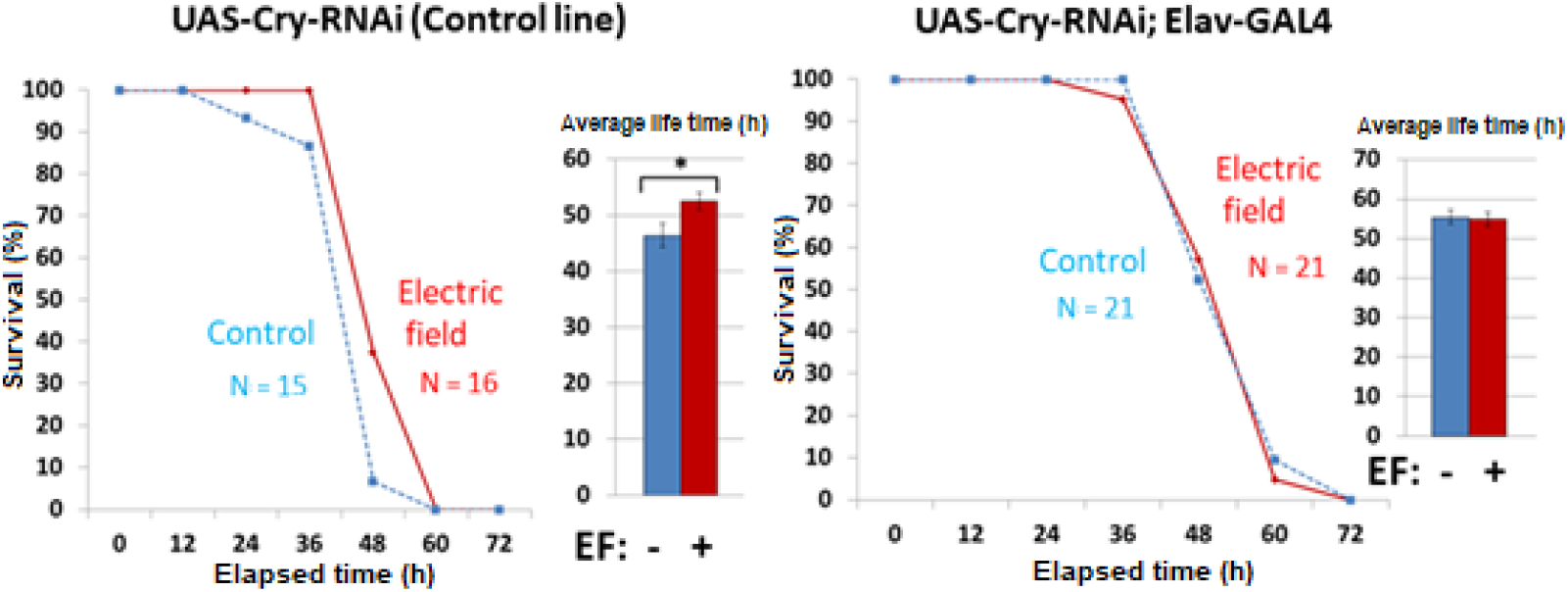
Effects of electric field on lifespan of wild and *Cry* gene mutant *Drosophila melanogaster* under starvation. a. The lifespan of wild flies with both of sex was ~20% prolonged by 50 Hz of EF exposure that started at 2–3 days after eclosion. Starvation was in 1% agarose (that maintained humidity). Red and blue, average lifespans with and without EF exposure. EF, electric field. b. Schematic of *Cry* gene mutants used to investigate *Cry* gene involvement in lifespan prolonged by EF. c. and d. Lifespan was not elongated in all three types of *Cry* mutants (c) or in flies with *Cry* RNAi (d). Exposure to EF increased lifespan of WT, but not of *Cry*-deficient (*Cry^b^*, *Cry^01^, Cry^03^* and *Cry* RNAi) lines. In contrast, the lifespan of control *Cry* RNAi parental strains was increased. EF, electric field, WT, wild type.

To confirm that *Cryptochrome* is involved in the lifespan elongation mechanism as observed in sleep, we prepared the *Cryptochrome* mutants, *Cry^b^*, *Cry^01^*, and *Cry^03^* (Fig. 3b). Interestingly, EF exposure did not alter the lifespan under starvation in any of these mutant lines (Fig. 3c).

We also assessed the EF effect on the lifespan of RNAi mutant flies under starvation (Fig. 3d). Although EF elongated the lifespan of the parental line, that of the RNAi *Cryptochrome* mutant flies did not differ between sham and EF exposure. These data indicated that *Cryptochrome* is involved in lifespan elongation caused by EF exposure.

### Metablome analysis showed increased ATP levels in flies

Although we showed that the *Cryptochrome* gene is involved in sleep and the lifespan of flies, the involved metabolic pathways remained unknown. We compared the metabolomes of WT and *Cry^b^* mutant flies with or without EF exposure to determine whether EF induces differences in metabolites. Figure 4a shows many differences between control and EF-exposed flies. The difference between Oregon R and *Cry^b^* mutant flies might be explained by the genetic backgrounds of the parent flies. The differences between control and WT exposed to EF were significant.

**Fig. 4.**
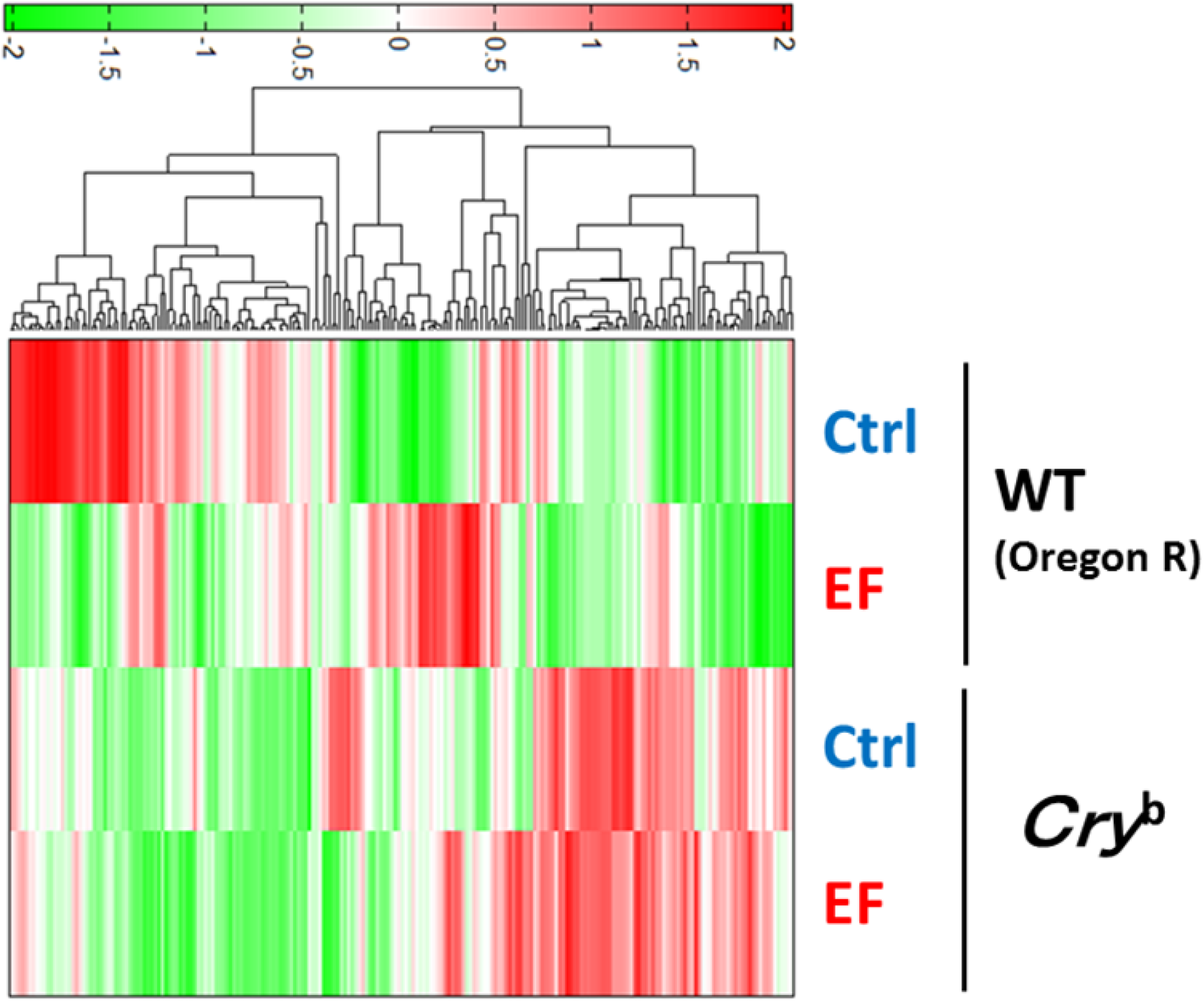
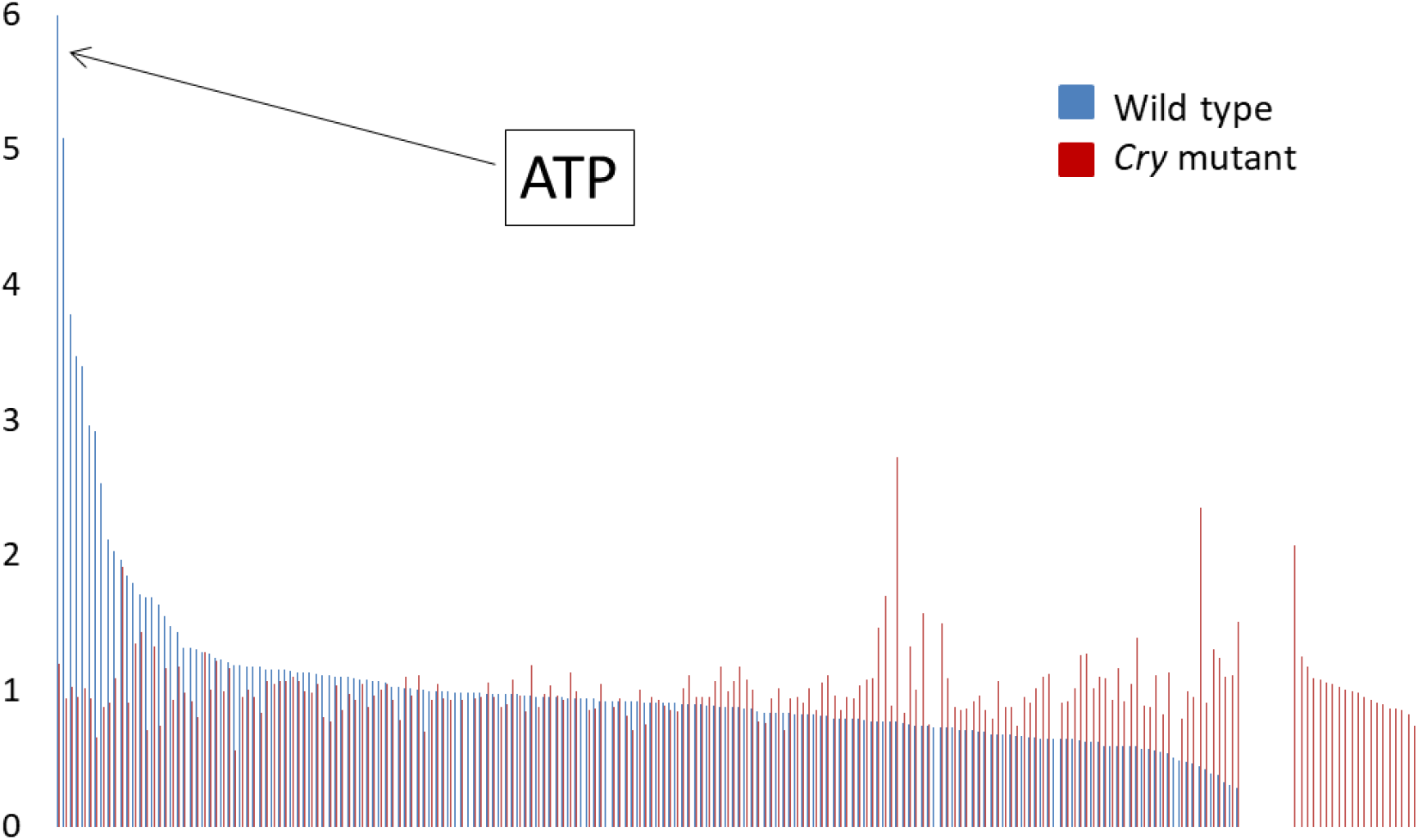
Metabolome analysis after EF exposure. a. Hierarchical cluster analysis of metabolome separation. Heat maps show differences in metabolites between control (Ctrl) and EF exposure. Red and green, high and low expression, respectively. Columns represent different conditions (Ctrl and EF, constant darkness without and with electric field. b. Comparison of upregulated compounds in WT and *Cry* mutant flies. Expression of compounds in WT and *Cry* mutant is arranged in descending order. ATP is most upregulated by EF. ATP, adenosine triphosphate; EF, electric field, WT, wild type.

Figure 4b shows an increased ratio of metabolites in EF-compared with sham-treated flies. The graph is arranged in decreased order for wild type flies (blue) and *Cry^b^* mutant flies (red). The difference between control and EF-exposed WT was significant compared with the *Cry^b^* mutant. Table 1 summarizes the data. The abundance of ATP was 5.99-fold higher in EF-than sham-treated flies, but that in *Cry^b^* mutant flies was ~1.2-fold higher (Fig. 4b, Table 1). These data indicated that EF exposure significantly increased ATP level and several nucleic acid metabolism in whole fly.

**Table 1.** Top 17 compounds upregulated more in WT than *Cry* mutant. Abundance of ATP is Ca. 5.0-fold higher in WT than *Cry* mutant.

|  |  | **Wild type** | ***Cry<sup>b</sup>*** |
| --- | --- | --- | --- |
| 1 | ATP | 5.99589 | 1.20585 |
| 2 | UDP | 5.09161 | 0.94851 |
| 3 | GTP | 3.79169 | 1.02959 |
| 4 | Trehalose 6-phosphate | 3.48234 | 0.95581 |
| 5 | ITP | 3.40563 | 1.01659 |
| 6 | Asp | 2.97017 | 0.94291 |
| 7 | O-Phosphoserine | 2.91937 | 0.65825 |
| 8 | CDP | 2.53999 | 0.88619 |
| 9 | PRPP | 2.11901 | 0.91622 |
| 10 | ADP | 2.0375 | 1.09812 |
| 11 | Myristoleic acid | 1.97591 | 1.91467 |
| 12 | UDP-glucose<br>UDP-galactose | 1.85463 | 0.91975 |
| 13 | 3-Phosphoglyceric acid | 1.80111 | 1.35414 |
| 14 | Fructose 1,6-diphosphate | 1.71542 | 1.43318 |
| 15 | Threonic acid | 1.69059 | 0.7116 |
| 16 | Dihydroxyacetone phosphate | 1.69036 | 1.32744 |
| 17 | Glutathione (GSSG)_divalent | 1.64224 | 0.74731 |

## Discussion

Exposure to 50 Hz of EF (35 kv/m, constant) during the daytime improved sleep quality in WT flies, whereas nighttime exposure did not. Interestingly, this effect did not observed in *Cry^b^* mutant flies. The fact that the number of sleep bouts decreased but the sleep bout length and total sleep time increased with daytime EF exposure suggests that sleep fragmentation was avoided and the duration of each bout of sleep was increased. This phenomenon is considered to be good marker of sleep quality and health in medical science. Exposure to 50Hz EF at night did not exert such effects. This difference might be explained by the circadian expression of *Cry* at different times (19) of the day.

We also found that wild type fly placed in 50Hz Electric Field elongated lifespan about 18%. Such lifespan elongation effect was not observed by using several lines including *Cry* gene mutant strains and *Cry* RNAi strain. CRYPTOCHROMEs (CRY) are structurally related to ultraviolet (UV)/blue-sensitive DNA repair enzymes called photolyases but lack the ability to repair pyrimidine dimers generated by UV exposure. The *Cry* gene was originally identified as a blue light photoreceptor in plants (20, 21). The role of CRY in *Drosophila* is cell-autonomous synchronization and the entrainment of circadian clocks (19). However, CRY plays a key role in magnetic field detection (8~14). Here we show that the putative magnetoreceptor protein and UV-A/blue light photoreceptor, CRYPTOCHROME is deeply involved in electric field receptor in animals. CRYPTOCHROMEs 1 and 2 (CRY1 and CRY2) have circadian clock functions and detect magnetic fields in various animals (22).

Metabolome comparisons between intact WT and *Cry* gene mutant strains after exposure to EF indicated that ATP was ~5-fold more abundant in the WT. Daytime exposure to a 50Hz EF might activate cellular activities or motility in flies because ATP is the “principal energy currency” in metabolism and the most versatile small molecular regulator of cellular activities (23).

The present findings provide the first genetic evidence of a CRY-based electric field sensitive system in animals.

